# Why do phylogenomic analyses of early animal evolution continue to disagree? Sites in different structural environments yield different answers

**DOI:** 10.1101/400465

**Authors:** Akanksha Pandey, Edward L. Braun

## Abstract

Phylogenomics has revolutionized the study of evolutionary relationships. However, genome-scale data have not been able to resolve all relationships in the tree of life. This could reflect the poor-fit of the models used to analyze heterogeneous datasets; that heterogeneity is likely to have many explanations. However, it seems reasonable to hypothesize that the different patterns of selection on proteins based on their structures might represent a source of heterogeneity. To test that hypothesis, we developed an efficient pipeline to divide phylogenomic datasets that comprise proteins into subsets based on secondary structure and relative solvent accessibility. We then tested whether amino acids in different structural environments had different signals for the deepest branches in the metazoan tree of life. Sites located in different structural environments did support distinct tree topologies. The most striking difference in phylogenetic signal reflected relative solvent accessibility; analyses of sites on the surface of proteins yielded a tree that placed ctenophores sister to all other animals whereas sites buried inside proteins yielded a tree with a sponge-ctenophore clade. These differences in phylogenetic signal were not ameliorated when we repeated our analyses using the site-heterogeneous CAT model, a mixture model that is often used for analyses of protein datasets. In fact, analyses using the CAT model actually resulted in rearrangements that are unlikely to represent evolutionary history. These results provide striking evidence that it will be necessary to achieve a better understanding the constraints due to protein structure to improve phylogenetic estimation.

## Introduction

The growing availability of very large datasets, including genome-scale datasets, has revolutionized the field of phylogenetics. Phylogenomics, the use of very large-scale molecular datasets to address phylogenetic problems, was suggested to be a way to “end incongruence” by reducing the error associated with analyses of small datasets (Gee 2003). Phylogenomics has had many successes (e.g., Rokas et al. 2003; Nishihara et al. 2007; Misof et al. 2014;Wickett et al. 2014) However, many relationships in the tree of life remain problematic, typically as a limited number of specific nodes within otherwise well-resolved phylogenomic trees (e.g., Nosenko et al. 2013; Cox et al. 2014; Jarvis et al. 2014; Whelan et al. 2015; Parks et al. 2018). Multiple distinct resolutions of those problematic relationships have emerged in different studies (e.g., Ryan et al. 2013; Moroz et al. 2014; Dunn et al. 2015; King and Rokas 2017; Simion et al. 2017), sometimes with strong support. These alternative resolutions of the tree can be viewed as distinct “signals” in the data; by signal we mean the topology that emerges when the data are analyzed using a specific method, regardless of whether that signal is historical or non-historical in nature. A major goal of phylogenomics is the identification of the historical signal (i.e., the species tree) amid the myriad other signals in genomic data (e.g., Castoe et al. 2009; Edwards 2009; Townsend et al. 2012; Salichos and Rokas 2013; Reddy et al. 2017). Rather than putting an end to incongruence, phylogenomics has actually served to highlight the complexity of the signals present in genomic data.

The idea that non-historical signals can overwhelm historical signal substantially predates the phylogenomic era (e.g., Felsenstein 1978; Hendy and Penny 1989). However, the availability of genome-scale data emphasized the large number of cases where that conflicting signals emerge in phylogenetic analysis. The most extreme cases correspond to those where analytical methods are subject to systematic error, where non-historical signal overwhelms historical signal. In those parts of parameter space, increasing the amount of data will cause phylogenetic methods to converge on inaccurate estimates of evolutionary history with high support. Thus, phylogenomic analyses should lead either to high support for the true tree (the desired result) or to high support for an incorrect tree (if the analytical method is subject to systematic error). Cases where support is limited despite the use of large-amounts of data could reflect one of two phenomena: 1) the data contains a mixture of signals, some historical and some that are misleading; or 2) that the underlying species tree contains a hard polytomy. Understanding the distribution of historical and non-historical signal(s) in large-scale data matrices could allow better understanding of analytical methods and their limitations with phylogenomic datasets.

The problem of systematic error in phylogenomic analyses has been addressed using two basic approaches: 1) by data filtering focused on removing the misleading data (e.g., Collins et al. 2005; Liu et al. 2015); and 2) by using improved, typically more complex and (presumably) more “biologically-realistic” models of sequence evolution (e.g., Steel 2005; Wilke et al. 2012). These approaches are interrelated. The goal of data filtering is the removal of data that violate model assumptions whereas the goal of conducting analyses using more biologically realistic models is to improve the fit of the model to the data. The fundamental assumption underlying both of these approaches is that phylogenomic datasets are heterogeneous; i.e., that a single, simple model is unlikely to provide an adequate to fit to the data. More complex models typically introduce biological realism by adding heterogeneity in the model, which could address this issue without requiring the exclusion of data. Even restricting to single type of data, such as proteins, a number of heterogeneous models have been proposed (e.g., CAT; Lartillot and Philippe 2004), Thorne-Goldman-Jones structural models (Thorne et al. 1996; Goldman et al. 1998), and structural mixture models (Le and Gascuel 2010), although the degree to which these approaches actually ameliorate the impact of misleading signals remains a subject of debate (Whelan and Halanych 2016). Moreover, both approaches ultimately require understanding the signals present in the data.

Examining conflicting signals could provide as way to determine whether those signals are associated with specific parts of the phylogenomic data matrices, and may provide insights into the biological basis for the heterogeneity. Subdividing data matrices into individual loci is unlikely to be informative because individual loci are short and therefore have limited power to resolve difficult nodes (Patel et al. 2013); moreover, individual loci can be associated with distinct gene trees due to factors such as the multispecies coalescent (Maddison 1997; Slowinski et al. 2007; Edwards 2009). However, it should be possible to examine signal as long as the total amount of data is large enough to overcome stochastic error (i.e., if a sufficient number of sites and gene trees are available). There are two alternative hypotheses regarding the distribution of signal in any phylogenomic dataset:

H_0_: Conflicting signals are randomly distributed with respect to functionally-defined subsets of the data. This hypothesis predicts that separate analyses of those functionally-defined subsets of the data matrix will yield trees with the same topology (probably with lower support than the analysis of the complete dataset due to the smaller size of the subsets and the fact that conflicts within the data subsets are still present).
H_A_: Conflicting signals are non-randomly distributed with respect to functionally-defined data subsets. Different subsets of the data matrix defined using functional information are associated with distinct signals (i.e., analyses of those subsets yield different topologies when the subsets are analyzed separately).

Obviously, these hypotheses can only be tested for specific definitions of data subsets; failure to reject H_0_ could reflect a genuinely random distribution of signal or it could reflect failure to define those data subsets in an appropriate manner. The data subsets should also be large enough to overcome stochastic error (although cases where the data subsets are reduced to the point where stochastic error dominates the analysis are likely to resolve difficult nodes randomly with low support). However, if H_A_ can be corroborated for a specific way of dividing phylogenetic data matrices into subsets using functional criteria (e.g., coding versus non-coding, highly transcriptionally-active versus largely untranscribed, and so forth; e.g., Reddy et al. 2017) it is likely provide substantial information about the patterns of sequence evolution (Wilke 2012). That information is likely to be useful for phylogenetic model development.

Coding sequences are often used to examine phylogenetic relationships deep in the tree of life (e.g., He et al. 2014; Wickett et al. 2014; Raymann et al. 2015) and they are good candidates for this type of signal exploration. There are many biologically-motivated ways to divide proteins into subsets that are likely to have been subjected to different selective pressures to maintain their structure and function. Any approach used to subdivide protein alignments into subsets for phylogenetic signal exploration must meet two criteria: 1) it should result in subsets that are stable over evolutionary times; and 2) practical ways to assign aligned sites to subsets exist. Protein structure provides an obvious way to divide phylogenomic data that meets both criteria. Protein structures diverge much more slowly than protein sequences (Lesk and Chothia 1986; Illergård et al. 2009), so it is reasonable to assume that alignments can simply be divided into structural classes that remain relatively constant over evolutionary time. Modern protein secondary structure prediction methods (e.g., Magnan and Baldi 2014) have proven to be very accurate (able to classify ∼90% of residues accurately for proteins with homologs in pdb) so it should also be possible to construct analytic pipelines that meets the second requirement. Obviously, it is possible that conflicting signals are randomly distributed with respect to any particular approach to subdividing proteins based on their structure, but there are a number of ways to subdivide protein structures that have been extensively studied. Therefore, it is possible to propose testable sub-hypotheses of H_A_ by focusing on strategies that reflect those extensively studied subdivisions. All that remains to test these hypotheses is a phylogenomic dataset likely to have internal conflicts; ideally, the test dataset would be large, analyses of the data should have fairly limited support for one or more focal clades.

We chose the early evolution of metazoa as a “model system” to examine whether different signals are associated with sites in different protein structural environments. There are several reasons why we expected the deepest branches in the animal tree of life to provide an ideal way to examine our hypotheses. First, the divergences among the major metazoan lineages are fairly ancient, so protein structure is likely to have had a fairly large impact on observed site patterns in the data. Second, deep metazoan phylogeny has been a difficult phylogenetic problem but there are a limited number of plausible trees (fig. 1). Specifically, the traditional hypothesis (fig. 1a), which also has support in some phylogenomic analyses (e.g., Philippe et al. 2009; Pick et al. 2010; Pisani et al. 2015; Simion et al. 2017), places sponges sister to all other metazoans. The alternatives are the hypothesis that ctenophores are sister to all other metazoans (fig. 1b), a hypothesis supported by many analyses of large-scale datasets (e.g., Dunn et al. 2008; Hejnol et al. 2009; Nosenko et al. 2013; Moroz et al. 2014; Whelan et al. 2015; Whelan et al. 2017). A third hypothesis with a ctenophore-sponge clade (fig. 1c); has only been found in the Ryan et al. (2013) genomic dataset (hereafter “RG”). There is a good argument for T3 being the least likely topology since it requires many of the same assumptions as T2 (e.g., it requires multiple origins of the nervous system or a single origin of the nervous system followed by a loss in sponges) but T3 has not emerged in most other large-scale analyses. Moreover, support for the ctenophore-sponge clade is limited in some analyses of the RG data and recovery of the clade is sensitive to taxon sampling, suggesting the RG dataset might have fairly strong internal conflicts. These internal conflicts could actually useful precisely because it should be possible to ask whether the conflicts are distributed randomly (H_0_) or non-randomly (H_A_) with respect to various subsets of the data defined using non-phylogenetic criteria. Herein, we subdivided the globular proteins in the RG dataset into subsets based on protein structure and explored the distribution of signal in different structural environments. Our results corroborated H_A_, revealing that the signal associated with solvent exposed residues differed from associated with buried residues. This has implications for the fit of models that are currently used for phylogenetic analyses of protein sequences.

**Fig. 1.**
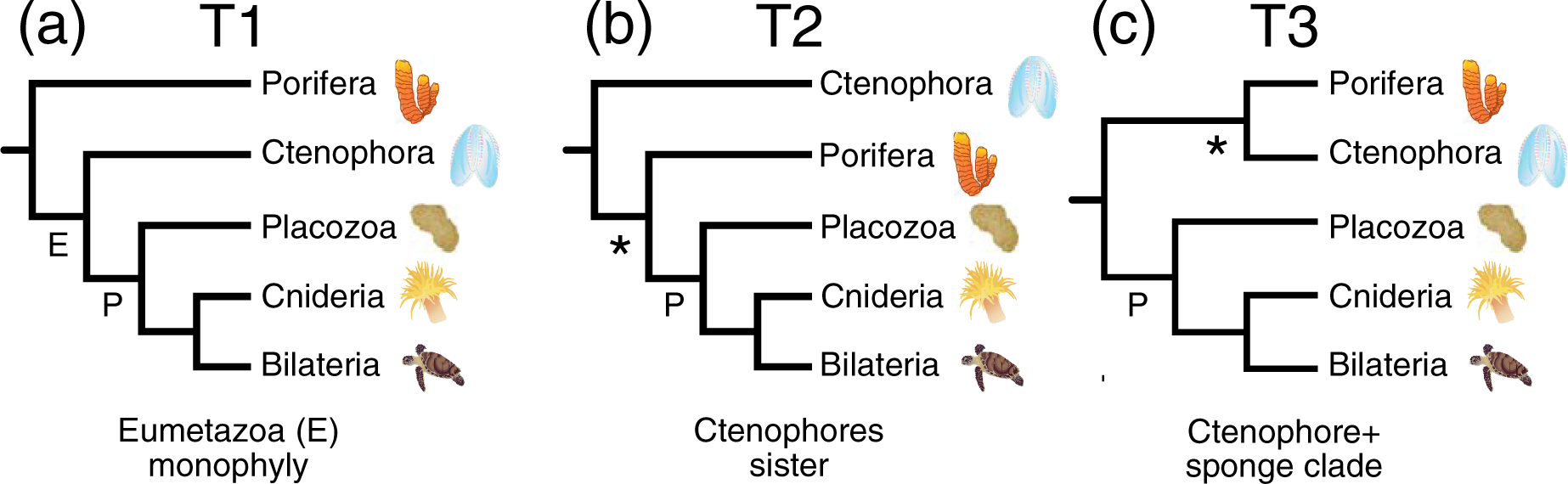
Topologies for the deepest branches in the metazoan tree recovered in phylogenomic analyses. (*A*) Porifera (sponges) sister to all other metazoa. This hypothesis includes a clade designated Eumetazoa (E). (*B*) Ctenophora (comb jellies) sister to all other metazoa. (*C*) A sponge-ctenophore clade sister to all other animals. All trees shown include a clade named Parahoxozoa (P) (Ryan et al. 2010). The topology for Parahoxozoa was fixed based on King and Rokas (2017).

## Results

### Sites in distinct structural environments have different signals

Different structural classes were associated with different phylogenetic signals based on analyses using standard empirical models. The most obvious difference was evident when we divided a dataset with 231 globular proteins [the “filtered Ryan genomic” (FRG) dataset, see Methods] using relative solvent accessibility. Analyses of solvent exposed sites placed ctenophores sister (T2) to all other metazoa whereas analyses of the buried sites recovered a sponge-ctenophore clade (T3) (fig. 3). In all cases, the best-fitting standard empirical model was LG (Le and Gascuel 2008) but our results were robust to the use of different models for ML analyses (table 1). In fact, they were robust to the use of the 20-state general time reversible (GTR) model, which had a better fit to the structurally-defined subsets of the FRG data than the LG model despite the large number of free-parameters that must be optimized for that model. Ultimately, we were unable to find a model that changed our conclusions.

**Table 1.** Log likelihood values and AIC*_c_* scores obtained using standard empirical models and GTR model optimized for exposed and buried site classes.

| **Structural Subset** | **Model** | **T2** | **T3** | **Cni+Bil<sup>a</sup>** | **Cni+Pla<sup>b</sup>** | **lnL** | **AIC<sub>c</sub>** |
| --- | --- | --- | --- | --- | --- | --- | --- |
| Exposed | GTR | 92 | - | - | 100 | -1212232.985 | **2424956.474** |
|  | LG | 100 | - | - | 100 | -1222489.017 | 2445088.162 |
|  | WAG | 87 | - | - | 100 | -1225898.553 | 2451907.235 |
|  | VT | 96 | - | - | 98 | -1225672.633 | 2451455.395 |
|  | rtREV | 90 | - | - | 98 | -1222733.929 | 2445577.986 |
|  | JTTDCMUT | 94 | - | - | 98 | -1229028.451 | 2458167.031 |
| Buried | GTR | - | 82 | 61 | - | -1045694.924 | **2091880.072** |
|  | LG | - | 87 | 62 | - | -1050022.577 | 2100155.267 |
|  | WAG | - | 85 | 58 | - | -1054829.256 | 2109768.626 |
|  | VT | - | 81 | 61 | - | -1059148.671 | 2118407.455 |
|  | rtREV | - | 89 | 52 | - | -1054432.123 | 2108974.359 |
|  | JTTDCMUT | - | 81 | - | 54 | -1061103.295 | 2122316.704 |

<sup>a</sup> Support for cnidaria + bilateria clade
<sup>b</sup> Support for cnidaria + placozoa clade

**Fig. 2.**
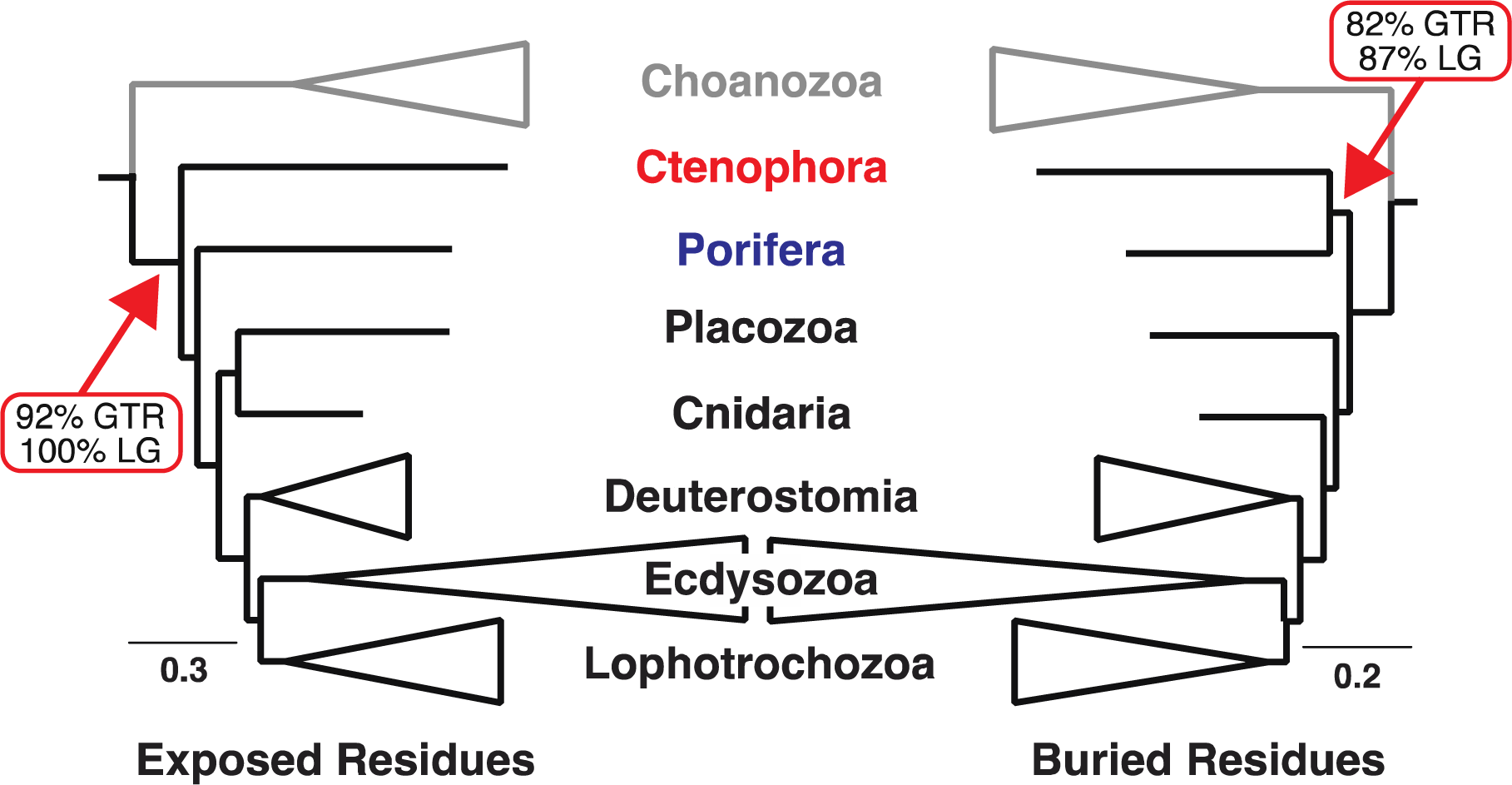
Analyses of sites from different structural environments reveal conflicting phylogenetic signals. We show simplified RAxML trees with both trees are limited to the metazoan ingroups and the choanozoan outgroup [i.e., only Apoikozoa *sensu* Budd and Jensen (2017) are shown]. The position of the root drawn in these trees was established by the outgroup taxa (the holozoans *Capsaspora* and *Sphaeroforma* and the fungi *Saccharomyces* and *Spizellomyces*). Bootstrap support for the positions of sponges and ctenophores given the GTR and LG models is indicated next to the arrow. All trees are available from Zenodo (Pandey and Braun 2018).

**Fig. 3.**
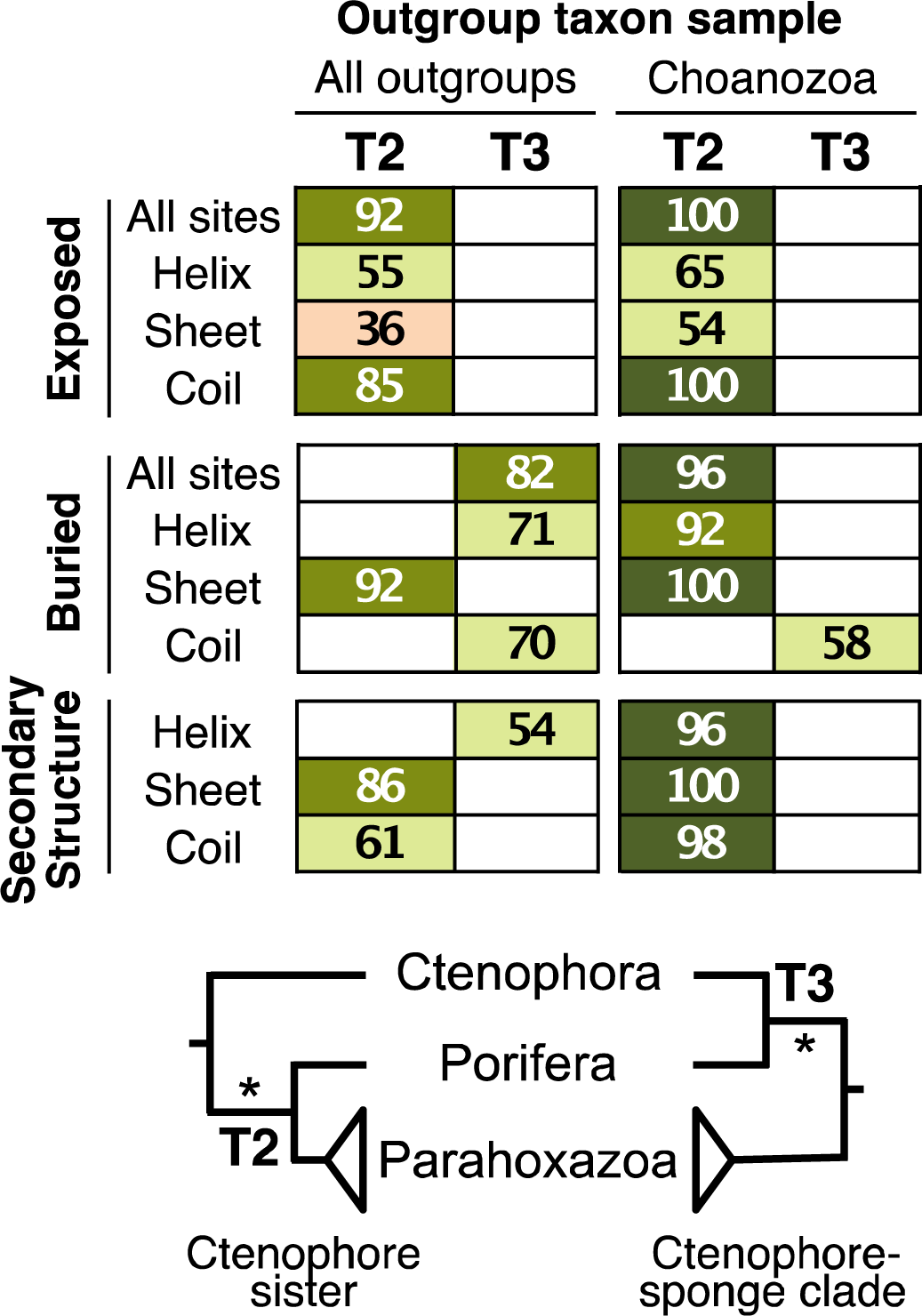
Heat map showing support for tree topologies obtained using various structural classes and taxon samples. In the online version colors indicate support values (Dark green > 95, lighter Dark green >75, Yellow > 50 and Pink < 50; No color: Tree not present)

Further subdivision of the FRG data based on other secondary structure information (helix, sheet, and coil) generally had less impact on topology, although we did find that buried sheet residues placed ctenophores sister to all other metazoans (unlike analyses of the other buried residues). However, all analyses of sheet residues, even when those sites were divided into solvent exposed versus buried sites, resulted in an unexpected clade comprising sponges and placozoa (all treefiles are available from in Pandey and Braun 2018). Subdividing the helix and coil sites into exposed versus buried subsets still revealed the different signals evident when the exposed and buried sites were defined globally (Pandey and Braun 2018). Overall, these results indicate that, if the topology at the base of metazoans is used as a way to measure signal, the data subsets with the largest difference in signal are the exposed versus buried sites and that, with respect to the deepest branches in the metazoan tree, the signals in the RG dataset support either tree T2 or T3 (fig. 2).

### Decisive sites reveal conflicts within each structural class

Strong phylogenetic signals are often limited to a subset of genes (Salichos and Rokas 2013; Brown and Thomson 2016; Shen et al. 2017) or even to specific sites within genes (Evans et al. 2010; Kimball et al. 2013). Examining these strong phylogenetic signals can provide a way to determine the amount of conflict within the data subsets. We examined the number of “decisive sites” (sites that strongly support either of two trees in fig. 2) in the exposed and buried classes. There were significant differences (*P* < 0.01, Fisher’s exact test) in the numbers of decisive sites that correspond to solvent exposed sites versus those associated with buried residues (table 2). Taken as a whole, these results indicate that differences in signal in different parts of the FRG dataset that can be detected in comparisons of solvent exposed versus buried sites does not reflect an unusual concentration of decisive sites in any particular gene within the FRG dataset; instead, the contrasting signals appear to be a more universal feature of analyses focused on sites in the two different structural environments.

**Table 2.**
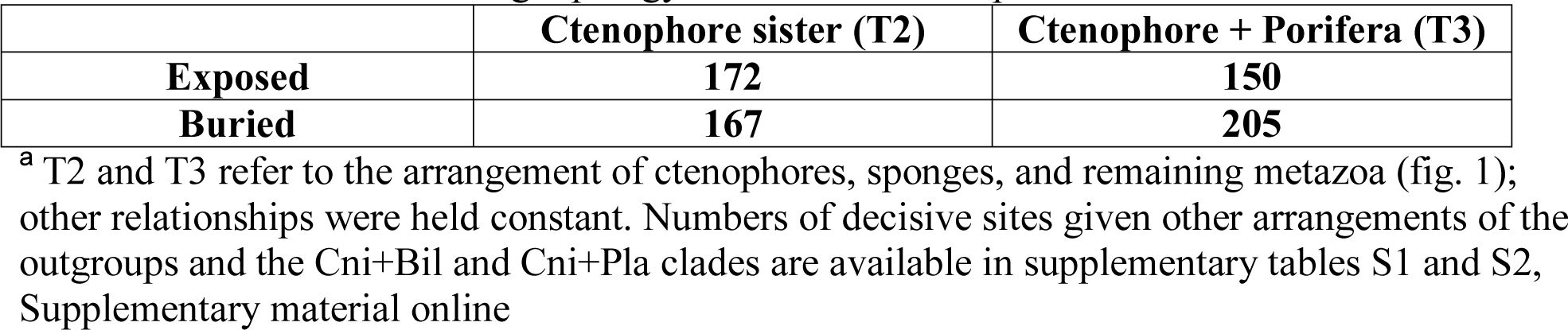
Decisive sites favoring topology T2 vs. T3 in the exposed and buried residues*^a^*.

### Reduced taxon sampling also reduces the differences in signal

Highly divergent outgroup taxa are known to affect phylogenetic inference (Philippe and Laurent 1998); to examine the impact of outgroup taxon sampling on the signal associated with various structural environments we removed four divergent outgroups [two fungi (*Saccharomyces* and *Spizellomyces*) and two taxa from the classes Mesomycetozoea (*Sphaeroforma*) and Filasterea (*Capsaspora*)] from the FRG dataset. This limits the outgroups to Choanozoa, the sister group of metazoa (Carr et al. 2008; Schalchian-Tabrizi et al. 2008). Analyses of exposed and buried residues using the FRG dataset with the reduced taxon sample converged on a single basal topology (ctenophores sister to all other metazoa) regardless of whether analyses used GTR with parameters optimized for each subset (fig. 3 and Pandey and Braun 2018) or standard empirical models (table 1). However, the position of cnidaria and *Trichloplax* varied in analyses using the reduced taxon sample (table 1 and table 3), underlining both the complexity of the signals in the FRG dataset and differences among the structural environments.

**Table 3.** Log likelihood and AIC*c* scores for various CAT model profile mixtures as implemented in IQ-TREE.

| CAT model | Outgroup | Dataset | T2 | T3 | Cni+Bil | Cni+Pla | Pori+Pla | lnL | $AIC_c$ |
| --- | --- | --- | --- | --- | --- | --- | --- | --- | --- |
| C10 | All outgroups (RG) | Exposed | 100 |  |  | 100 |  | -1225103.471 | 2450339.126 |
| C20 |  | Exposed |  | 63 |  | 100 |  | -1216444.36 | 2433040.964 |
| C30 |  | Exposed | 100 |  |  |  | 72 | -1214319.593 | 2428811.499 |
| C40 |  | Exposed |  | 100 |  | 96 |  | -1212974.829 | 2426142.047 |
| C50 |  | Exposed | 100 |  |  |  | 73 | -1212360.091 | 2424932.657 |
| C60 |  | Exposed |  | 100 |  | 93 |  | -1211470.44 | **2423173.448** |
| C10 |  | Buried |  | 83 | 78 |  |  | -1053873.212 | 2107878.588 |
| C20 |  | Buried |  | 97 | 80 |  |  | -1048007.721 | 2096167.66 |
| C30 |  | Buried |  | 94 | 73 |  |  | -1046099.005 | 2092370.288 |
| C40 |  | Buried |  | 92 | 83 |  |  | -1044155.469 | 2088503.283 |
| C50 |  | Buried |  | 93 | 86 |  |  | -1043172.44 | 2086557.3 |
| C60 |  | Buried |  | 93 | 89 |  |  | -1042207.261 | **2084647.025** |
| C10 | Choanozoa only | Exposed | 100 |  |  | 100 |  | -964491.9951 | 1929100.133 |
| C20 |  | Exposed | 98 |  |  | 74 |  | -957706.2984 | 1915548.793 |
| C30 |  | Exposed | 96 |  |  | 69 |  | -956161.8764 | 1912480.01 |
| C40 |  | Exposed | 95 |  |  | 71 |  | -955297.444 | 1910771.215 |
| C50 |  | Exposed | 92 |  |  | 51 |  | -953519.0992 | 1907234.604 |
| C60 |  | Exposed | 90 |  | 100 |  |  | -952582.9203 | **1905382.332** |
| C10 |  | Buried |  | 50 | 46 |  |  | -837871.4589 | 1675859.045 |
| C20 |  | Buried |  | 37 | 34 |  |  | -833463.787 | 1667063.748 |
| C30 |  | Buried |  | 75 | 63 |  |  | -832000.2662 | 1664156.761 |
| C40 |  | Buried |  | 66 | 63 |  |  | -830714.1456 | 1661604.582 |
| C50 |  | Buried |  | 45 | 45 |  |  | -829970.749 | 1660137.858 |
| C60 |  | Buried |  | 74 | 79 |  |  | -829121.6382 | **1658459.713** |

### Sites in different structural environments exhibit distinct patterns of sequence evolution

The two different topological signals evident in analyses of datasets that contain only exposed or buried residues (fig. 2) emerged regardless of whether analyses were conducted using standard empirical models (e.g., LG) or the GTR model optimized on the FRG dataset as a whole (rate matrix parameter estimates are available from Zenodo; Pandey and Braun 2018). Thus, the observed signal variation in exposed and buried classes cannot simply reflect the poor fit of standard empirical models to FRG, since the effect persists when the model is optimized for the dataset. However, standard empirical models and GTR optimized for a specific large-scale protein dataset share the property that the rate matrices reflect the patterns of sequence evolution for a mixture of sites from all structural classes. Thus, the values in the empirical rate matrix probably reflect compromise values relative to the values that would be estimated using structurally divided data. This led us to examine the degree to which estimates of GTR model parameters differ among sites in different structural environments.

A fundamental prediction of the hypothesis that distinct signals will emerge in analyses of sequence alignments comprising only those sites associated with a specific structural partition is that patterns of sequence evolution in each structural environment differ. If so, differences in the patterns of sequence evolution should be evident in estimates of GTR matrix parameters for each structural subset and these estimates should be fairly different from the values for standard empirical models. The model parameters for each structural environment are not only expected to differ from standard empirical models but also from each other. When the differences among models were examined by multidimensional scaling the strongest separation among models is related to the best models for the two solvent accessibility classes (fig. 4). Standard empirical models, like the LG, WAG, and Dayhoff models, formed a cluster closer to models estimated using buried residues (fig. 4 and supplementary figure S1, Supplementary Material online).

**Fig. 4.**
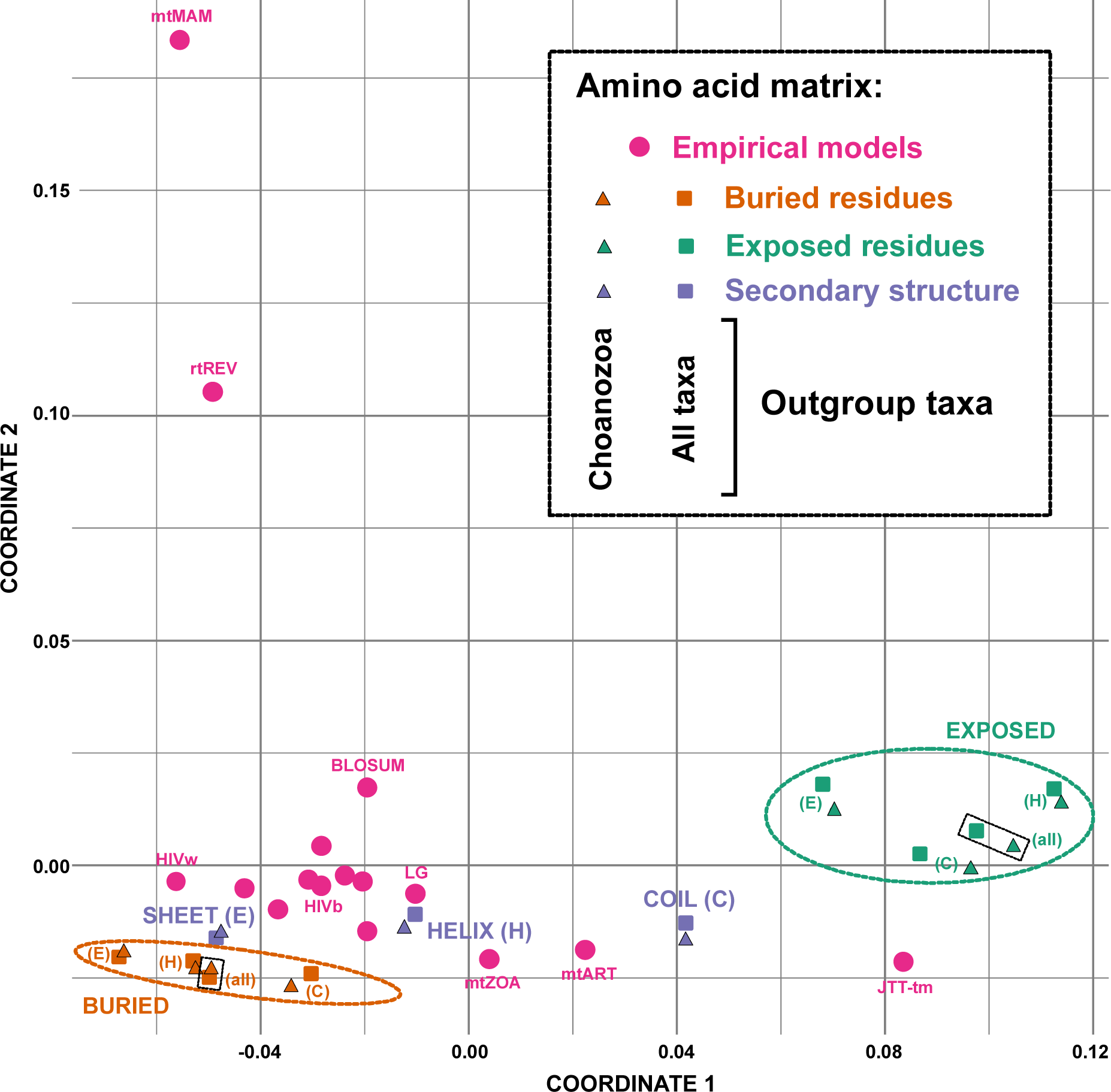
Multidimensional scaling plot showing the Euclidean distances between various amino acids exchange rate matrices. Different colors indicate different categories of the matrices in the online version (Green: exposed residues, Orange: buried residues, Purple: secondary structure, and Pink: standard empirical models).

The various structural subsets have different numbers of sites and the GTR model for amino acids has a large number of free parameters, raising the question of whether the observed differences in exchange rate parameter estimates simply reflect sampling error. The fact that the models appear to cluster in a manner that is correlated with structural class in multidimensional scaling space (e.g., note that buried sites and solvent exposed sites cluster in fig. 4 even when they are further subdivided into independent sets of helical, sheet, and coil sites) suggests that sampling error is unlikely to explain the observed difference. Nevertheless, we wanted to conduct an additional test focused on the impact of sampling variance. To do this we sampled sites from the FRG dataset randomly (i.e., without respect to structure) to generate datasets comparable in size to the structurally defined subsets and estimated model parameters on these random samples. Since the sites were sampled randomly, estimates of GTR model parameters should converge on the values estimated from the dataset as a whole, assuming that the number of sites that were sampled is sufficient to overcome sampling variance. We assessed the distance between the GTR model parameter estimates for the complete FRG dataset to those of randomly selected sites. We found that this distance rapidly decreased as the number of sites in the random sample increased (supplementary fig. S2, Supplementary Material online); the distances between the “global” model based on the complete FRG dataset and the estimates based on each structural subset are much greater than the distances expected based on sampling variance. This demonstrates that the differences in parameter estimates of structural classes differ much more than expected based on sampling variance.

### Site-heterogeneous profile mixture models can yield surprising topological changes

Site heterogeneous models, like the CAT model (Lartillot and Philippe 2004), represent another way to accommodate heterogeneity in the evolutionary process. CAT-type models assume aligned sites are drawn from a mixture with many distinct evolutionary processes (substitutional profiles) that differ in the equilibrium frequencies of the 20 amino acids. To complement the ML analyses using empirical models and the GTR model we analyzed the exposed and buried classes using the ML version of the CAT model (Le et al. 2008) with various numbers of profile mixture classes (C10-C60). We analyzed the full taxon set as well as reduced taxon set using the six ML variants of the CAT model implemented in IQ-TREE (Nguyen et al. 2015). If variation among sites in their propensity to accept specific amino acids is necessary to obtain accurate estimates of phylogeny then trees based on the two structural subsets (exposed versus buried) are expected to converge on the same topology.

Analyses of the exposed and buried sites from the full taxon set using the CAT model with various number of categories did converged on a single tree T3, with three exceptions (C10, C30 and C50 with exposed residues), but among C10, C30 and C50 two of these analyses converged on a tree with a sponge-ctenophore clade (table 3). In contrast, analyses of the reduced taxon sample using CAT models resulted in different topologies for exposed and buried classes (T2 and T3 respectively; table 3). Surprisingly, we found that analyses of buried residues yielded a non-monophyletic Deuterostomia; all profile mixtures placed the echinoderm in the FRG dataset (the sea urchin *Strongylocentrotus purpuratus*) sister to a clade comprising chordates and all of the protostomes in the taxon sample (fig 5). Furthermore, we noted that the mixtures with a larger number of category profiles mixtures (C40, C50 and C60) had a zero or near zero weights for the mixture components (parameter estimates are available from Zenodo; Pandey and Braun 2018). To test whether any of the observed topologies reflected differences between IQ-TREE and RAxML (e.g., in their search or numerical optimization routines) we confirmed that analyses using the site-homogeneous GTR model in IQ-TREE yielded the same results as analyses in RAxML (Pandey and Braun 2018). Although the most complex CAT model did have the best fit to the data based on the AIC*_c_* other results that emerged when we used CAT models, like the very low estimates of weights for specific substitutional profiles, could indicate over fitting of the model. Regardless of the AIC*_c_* values or parameter estimates, we emphasize two purely empirical observations that emerged when we used the ML variants of the CAT models: 1) the models were unable to increase congruence between estimates of phylogeny based on the exposed and buried residues in the FRG dataset; and 2) use of the CAT models revealed an unexpected signal in the buried residues (deuterostome non-monophyly) that is very likely to be non-historical.

**Fig. 5.**
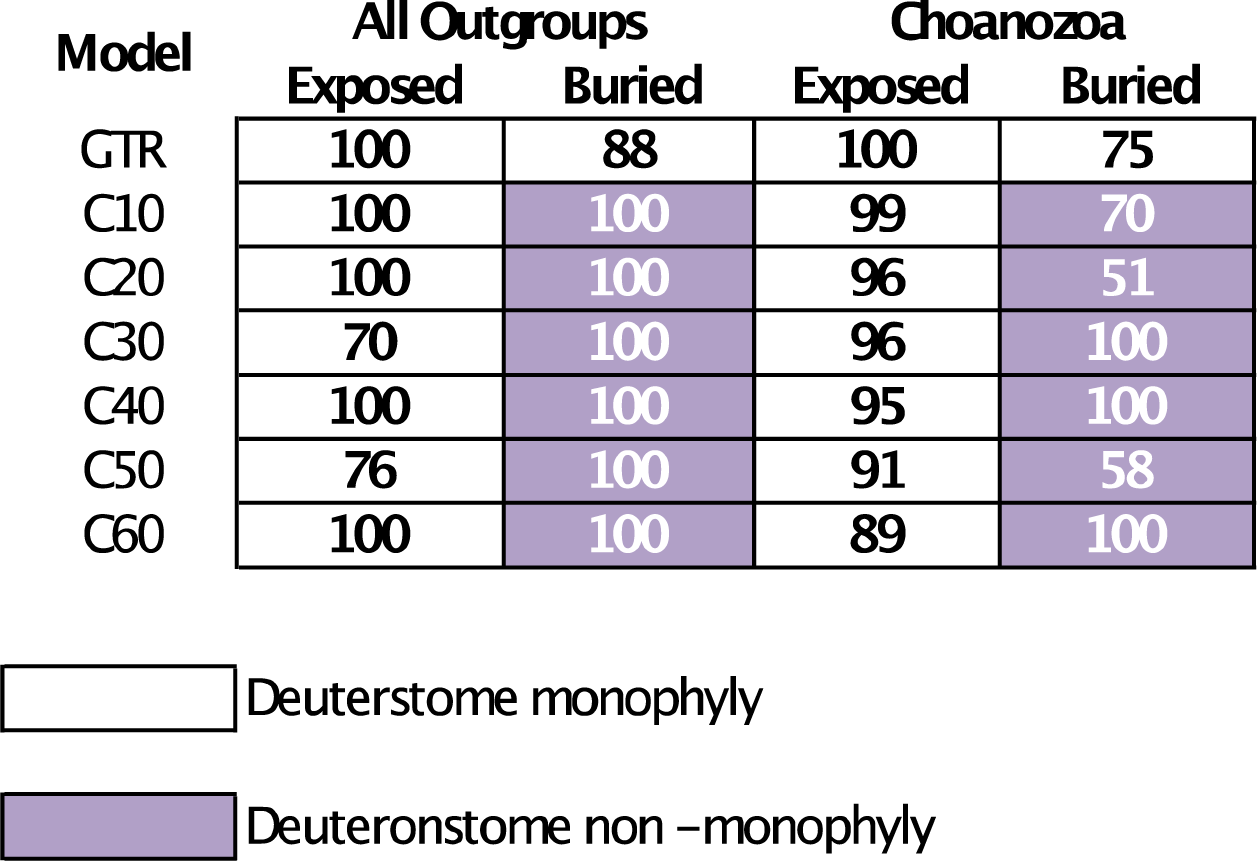
Heat map showing support for deuterostome monophyly for exposed and buried residues using GTR and CAT models. Colors in the online version indicate presence and absence of deuterostome monophyly (No color: present, Purple: absent).

## Discussion

Analysis of the FRG dataset with various structural classes resulted in different tree topologies (e.g., fig. 2) and there were significant differences in the numbers of decisive sites favoring distinct placements of the metazoan root (table 2). Both of these results strongly corroborate the H_A_, the hypothesis that conflicting phylogenetic signals are non-randomly distributed in parts of the dataset that can be defined using protein structure. Dividing protein alignments into subsets based on relative solvent accessibility (exposed sites versus buried sites) showed this difference in signal most prominently. The different signals in exposed and buried sites were evident regardless of whether the phylogenetic inference was conducted using standard empirical models or with GTR model parameters optimized on each structural subset. Other division strategies (i.e., dividing aligned sites based on secondary structure of a combination of secondary structure and relative solvent accessibility) also revealed some differences in signal, but they were not as strong as the differences between solvent exposed versus buried sites.

Limiting our analyses to Apoikozoa (the clade comprising Choanozoa and Metazoa; Budd and Jensen 2017) resulted in greater congruence between estimates of phylogeny based on exposed versus buried sites (fig. 3). With the exception of analyses based on the buried coil sites, all analyses conducted after excluding non-choanozoan outgroups supported the T2 (ctenophores sister; fig. 3). It is tempting to attribute these results to the suppression of long-branch attraction (LBA) since divergent outgroups were excluded. However, the major problems attributed to LBA at the base of metazoa has been attraction of the long ctenophore branch to the outgroups (e.g., Philippe et al. 2009; Nosenko et al. 2013); an corollary of that hypothesis that is implicit in most studies is that the true tree is T1 (sponges sister to other metazoa) and that recovery of T2 (ctenophores sister to other metazoa) reflects the impact of LBA. T3 (the ctenophore-sponge clade) is unlikely to reflect LBA, since the long ctenophore branch is united with the shorter branch leading to the sponge in that tree (supplementary fig. S3, Supplementary Material online). If T1 is hypothesized to be the true tree one also has to postulate that removing non-choanozoan outgroups caused analyses that support for one artifactual tree (T3) to shift towards support for another artifactual tree (T2) where the long ctenophore branch joins the outgroup branch. This would be surprising if LBA is the primary source of the systematic error. Hypothesizing that T2 is the true tree eliminates this paradox; all that one needs to postulate is that a change in taxon sampling that increases congruence between the partitions also reflects a shift toward the true tree in analyses of the buried sites. The basis for the increased congruence is unclear because estimates of GTR model parameters for each structural subset were fairly similar regardless of whether the complete taxon sample or the Apoikozoan taxon sample was used to estimate their values (fig. 4 and supplementary fig. 1, Supplementary Material online). All of these analyses emphasize that different signals, probably reflecting different patterns of sequence evolution, emerge in different structural environments.

### Different models for different structural environments

The obvious explanation for the conflicting signals in each structural class, especially the strong difference between the exposed and buried signals, is that the sites in each of these classes exhibit different patterns of sequence evolution. The poor fit of standard empirical models to the data could then result in an incorrect inference. This could result in the observed difference in signal, with one structural subset yielding the correct tree while analyses of the other one result an incorrect estimate of phylogeny. Alternatively, it is possible that analyses of both exposed and buried sites yield different topologies, both of which are inaccurate. GTR model parameter estimates for each class certainly indicate that the patterns of sequence evolution are different in each structural environment (fig. 4 and supplementary fig. S1, Supplementary Material online). However, conducting analyses using GTR model parameters optimized on each class did not cause analyses in the two classes to converge on a single topology (in fact, the topologies were unchanged relative to those that emerged in the analyses that used standard empirical models). Finally, we note one intriguing aspect of our model comparisons: the rate matrix parameters for most empirical models lie closer to the models based on buried sites (this was true for all empirical models trained on diverse datasets; i.e., all empirical models except those trained using the more limited organellar or viral datasets).

The basis for the differences we observed among the structural classes in their patterns of evolution almost certainly reflects differences in the nature of the purifying selection on sites in different structural environments. It is well known that protein evolution is heterogeneous and depends on the site-specific biochemical constraints like structure, dynamics, and biochemical functions (Echave et al. 2016). More precisely, depending on their position and role in the overall conformation and function of the protein specific sites will only accept subsets of the 20 amino acids, with all other possibilities being selected against. For example, if we consider the case of relative solvent accessibility, the sites that are exposed to an aqueous environment will include more polar residues whereas buried sites which form the cores of proteins and are inaccessible to solvent, will include more non-polar residues and be more resistant to changes in side chain volume (Gerstein et al. 1994). Similar differences among sites located in the major secondary structural classes have also been appreciated for some time (Thorne et al. 1996; Goldman et al. 1998; Le and Gascuel 2010), although the differences among the secondary structure classes does not appear to be as extreme as those based on solvent accessibility. Our observations suggest that these differences in evolutionary patterns among the structural classes can lead to different signals in each structural class.

Models of sequence evolution that incorporate protein structure have also been proposed in the context of phylogenetics. For example, Goldman et al. (1998) developed a hidden Markov model approach that assigns distinct rate matrices to sites based on in secondary structure and solvent accessibility (where secondary structure and solvent accessibility are the hidden states). Le and Gascuel (2010) implemented a mixture model that uses available protein structure annotations. Both of these studies revealed that efforts to acknowledge the different structural classes for sites in protein multiple sequence alignments can have a major impact on model fit, measured using the improvement in log likelihood values. However, both of those methods estimate a tree topology for the complete alignment; our approach of dividing proteins into structural classes follows the same basic idea but shifts the focus toward study the phylogenetic signals in these structural classes separately. Combined analysis of data is very useful since it has long been thought that best estimate should emerge from analyses that include as much evidence as possible (cf. Kluge 1989). However, the combined analysis paradigm suffers from the drawback that the phylogenetic analysis can simply converge reveal the topology that reflects the dominant signal in the data. Specifically, one might imagine a case similar to that illustrated by this study, where different structural classes are associated with distinct phylogenetic signals. If one further imagines observing one topology when the complete dataset is analyzed using a model that does not consider structure and a different topology when it is analyzed using second model that does consider structure one is faced with a question: do the observed topological differences reflect increased congruence between the structural classes (i.e., the tree topology signal associated with site patterns in each structural class disagrees given standard approaches but agrees when the data are analyzed using a structure-aware model) or do they reflect the strengthening (in relative terms) of the signal associated with one subset of the data without any increase in the agreement between the structurally-defined data subsets? As the true tree is unknown in all empirical studies, it is difficult to understand whether resulting topology reflects the true historic signal or a misleading (and potentially dominant) signal in the dataset. By dividing the data into subsets based on protein structure we can actually determine the signal in each of the structural classes actually changes to increase the congruence among the subsets.

### Site-heterogeneous models do not increase congruence in signal for different structural classes

Site-heterogeneous models like CAT has been used in many studies focused on the deep branches in the metazoan tree (e.g., Philippe et al. 2009; Pick et al. 2010; Philippe et al. 2011a; Egger et al. 2015) and it has been asserted that the CAT model is more realistic than empirical models or the GTR model (Lartillot and Philippe 2004; Liu et al. 2009; Tsagkogeorga et al. 2009; Finet et al. 2010; Philippe et al. 2011a; Nosenko et al. 2013). Analyses using CAT models have been suggested to be less prone to systematic error, such as LBA, than site-homogeneous models like the standard empirical models when they are applied to heterogeneous data (Lartillot et al. 2007; Brinkmann and Philippe 2008; Philippe et al. 2011b; Roure et al. 2013). If there is heterogeneity within protein secondary structural elements the CAT model could represent a better way to introduce heterogeneity into phylogenetic analyses than structure aware models (Thorne et al. 1996; Goldman et al. 1998; Liò et al. 1998). Nonetheless, it is important to recognize that all the models are approximations; whether or not any specific approximating model (e.g. the CAT model) can reveal the true historical signal in a specific dataset better than another approximating model (e.g., the GTR model) ultimately represents an empirical question.

We observed several indications that analyses of the FRG dataset using the CAT model did not behave ideally. If the CAT model captured the heterogeneity in each subset of the data and improved the results of phylogenetic analyses, then we would expect the trees estimated using the exposed and buried subsets of the data to converge on the same topology. We would also expect the resulting topology to be T1 or T2, since there are convincing arguments that T3 is the least plausible topology. However, our analysis using CAT models with various profile mixtures for exposed and buried residues either shifted back and forth between T2 and T3 or converged towards less plausible T3 topology. We believe that this indicates that CAT models are unable to capture the heterogeneity in these subsets in an appropriate manner. Furthermore, an unexpected clade uniting chordates and protostomes (i.e., the branch that implies deuterostome non-monophyly) also emerged in analysis using buried sites. We believe the observed support for the unexpected chordate-protostome clade as well as the shifts between T2 and T3 when increasing the number of categories provides evidence that CAT models are inappropriate for analyses of these data, possibly reflecting the overestimation of the number substitutional categories appropriate for larger datasets; the zero or near zero estimated weights for specific substitutional profiles provides another line of evidence for this idea. Regardless, our results certainly indicate that CAT models do not represent a panacea for phylogenetic analyses of proteins.

## Conclusions

We believe that our signal exploration focused on structure-based classes provided two valuable insights on how current phylogenetic methods deal with phylogenomic datasets. First, we corroborated the hypothesis that different phylogenetic signals are associated with specific parts of proteins that can be defined using non-phylogenetic criteria (in this case, structural criteria). Analyses of data from two structural environments resulted in conflicting trees that each had relatively high support, suggesting that this could reflect systematic error. Second, we provided evidence that one should not eschew the use of site-homogeneous models (like GTR or empirical models such as LG or rtREV) in favor of more complex site-heterogeneous models like CAT despite the better fit of the latter models based on commonly-used model selection criteria like the AIC*_c_*. It is important to recognize the difficulty to assess the fit of models in absolute terms (Gatesy 2007). Even the most complex models currently available for phylogenetic analyses, including the site-heterogeneous models, may be quite far from the true underlying processes of molecular evolution and, therefore, every bit as subject to systematic error as simpler models. Indeed, Sanderson and Kim (2000) worried that the very large AIC increases that accompany the addition of relatively small numbers of free parameters to phylogenetic models might indicate that all models that are being considered may be very far from the unknown true model. Perhaps the AIC*_c_* increases we observed indicates a case where the CAT model is indeed closer to the true model but deviates from that model in ways that obscure historical signal rather than revealing that signal. Although available site-heterogeneous models might be very useful in some contexts; they do introduce a large number of free parameters that may not be constrained in a biologically-realistic manner. Steel (2005) worried that very parameter-rich models would be impractical because they would have to have enough parameters “to fit an elephant” (based on the colorful metaphor that some have attributed to John von Neumann; Dyson 2004). The observed association between protein structure and phylogenetic signal points towards a way to overcome this “elephant factor”. Although it is probably impossible to devise a reasonably realistic model for any biological process that has a very low dimension, further analyses of protein evolution in the context of protein structure and function may allow us to constrain models of sequence evolution in ways that permit the true historical signal to be recovered more accurately, thereby addressing the shortcomings of available methods.

## Materials and Methods

### Dataset

The Ryan et al. (2013) genomic (RG) dataset comprises 242 orthologous protein-coding genes extracted from genomic data for 19 taxa. We used the alignments from Ryan et al. (2013), who aligned the sequences using CLUSTALW (Thompson et al. 2002) with default parameters and excluded poorly aligned regions using Gblocks (Castresana 2000). As a check, we visually inspected all of the alignments in Geneoius v. 9.1.5 (Biomatters Ltd., Auckland, New Zealand) and none of the Ryan et al. (2013) alignments appeared problematic. We used the TopCons prediction server (Tsirigos et al. 2015) to determine whether there were any transmembrane proteins in the RG data; we identified 10 transmembrane proteins (Pandey and Braun 2018). We removed the transmembrane proteins because our structural assignment pipeline is not appropriate for those proteins.

The 232-protein dataset that resulted when the transmembrane proteins were excluded had 102,000 sites and 19.81% missing data. We conducted preliminary analyses of this 232-protein dataset by dividing the alignment into structural subsets; analyses of the exposed and buried subsets resulted in two distinct trees identical to those in fig. 2. We then determined whether any specific gene strongly favored either topology using a “decisive gene” analysis (see below in the “phylogenetic analyses” section). We found that gene 41 had a strong signal favoring the buried topology. In an unpartitioned analyses using the LG model the likelihood difference given the two trees in fig. 2 was more than three-fold greater for gene product 41 than for any of the other proteins in the data matrix (Δ*ln*L = 106.63 favoring the buried tree for protein 41 compared to a range of Δ*ln*L = 9.39 to 28.67 for the other proteins; see Pandey and Braun 2018). Because protein 41 was an outlier relative to the other sequences (there are many reasons why it might be unusual, including a mistaken orthology call) we removed that gene to yield a 231-protein data matrix that we called the filtered Ryan genomic (FRG) dataset. We emphasize that removing protein 41 did not alter the trees recovered after dividing sites into subsets based on structure (see Results); however, we felt it was important to remove any outlier genes that might not be true orthologs.

We conducted BLASTP searches of UNIPROT (Uniprot Consortium 2018) to determine the identity and function of each gene (using annotation of the human sequence for most genes). There were 22 cases where the human sequence was absent from the RG alignment; in those cases, we used the *Drosophila* sequence to identify the genes. These functional annotations are available from Zenodo (Pandey and Braun 2018). This analysis revealed that remaining proteins represent a diverse set of globular proteins without an overrepresentation of sequences from a particular protein family making it a relatively unbiased dataset. We provide all protein alignments as Nexus files with structural annotation, generated as described below, in Zenodo (Pandey and Braun 2018). We analyzed two taxon samples: 1) the full taxon sample comprising all 19 taxa in the RG dataset; and 2) a reduced (apoikozoan) taxon sample that excluded four relatively divergent (non-choanozoan) outgroups (*Capsaspora, Saccharomyces, Sphaeroforma*, and *Spizellomyces*).

### Structural Class Assignment

We assigned secondary structures to the MSAs using the SSpro and ACCpro programs in the SCRATCH 1D suite (Cheng et al. 2005). SSpro classifies sequences into three secondary structural classes (helix, sheet, and coil) with a prediction accuracy of 92% for proteins with homologs in PDB (Magnan and Baldi 2014). ACCpro assigns each residue to one of the two categories: exposed (e) or buried (-) (Pollastri et al. 2002) with the latter defined as amino acids with <25% relative solvent accessibility; it has a predication accuracy of 90% (Magnan and Baldi 2014). We used a weighted consensus sequence representing each protein in the RG dataset as input for SSpro and ACCpro. We generated a weighted consensus sequence for each protein using the Henikoff and Henikoff (1994) method; the amino acid residue with the highest weight at each position was used in the consensus sequence. The structural data were extrapolated from the consensus sequence to the whole alignment and written as CHARSETS in the nexus file. We then extracted sites of a given structural class from all the genes and created a concatenated alignment for a given structural class. The perl program for this analysis is available from github (https://github.com/aakanksha12/Structural_class_assignment_pipeline).

### Phylogenetic Analyses

We used RAxML v. 8.2.4 for tree searches and log likelihood estimation. We examined a set of standard empirical models (LG, WAG, VT, JTT-DCMUT and rtREV) with ML estimation of amino acid frequency parameters (-PROTGAMMALGX option in RAxML) and we used GTR model parameters optimized on the complete dataset (the “grand GTR parameters”) and various subsets of the data (see below for a description of the structural partitions). In all cases we also used GTRGAMMA (i.e., the GTR model combined with a four-category discrete approximation to the Γ distribution that describes rates across sites). We assessed the nodal support using rapid bootstrap (Stamatakis et al. 2008) using the bootstopping criterion (Pattengale et al. 2009) as implemented in RAxML (i.e., the -N autoMR option).

To examine whether there were any observed differences among partitions in their signal we searched for decisive genes and decisive sites (cf. Kimball et al. 2013; Shen et al. 2017). Decisive genes and decisive sites are defined as those genes or sites with a very large impact on the likelihood given each a specific topology. We identified decisive genes and decisive sites by calculating per site likelihoods for each candidate topology using RAxML (via the ‘-f G’ option). To identify decisive genes we calculated the sum of the likelihood values within each gene and then calculated Δln*L* (ln*L* given tree 1 minus ln*L* given tree 2). At the level of individual sites we calculated Δln*L* for each site and, based on Kimball et al. (2013), viewed sites with Δln*L* > 5 standard deviations as decisive sites.

### Model estimation

We obtained ML estimates of the model parameters (amino acid exchangeabilities and amino acid frequencies) for each structural class using RAxML (Stamatakis 2014). Our approach relaxes the notion that all sites evolve following the same Markovian process, but it still assumes that all the sites in the same subset of the MSA follow the same stationary and homogeneous Markov process (i.e., each subset has its own set of GTR parameter estimates). We examined the following classes:

a. Two relative solvent accessibility (RSA) based classes (EXPOSED and BURIED).
b. Three secondary structure-based classes (HELIX, SHEET, and COIL).
c. Six classes, combining RSA and secondary structure (HELIX_EXP, HELIX_BUR, SHEET_EXP, SHEET_BUR, COIL_EXP, and COIL_BUR).

We estimated the GTR model parameters (-f e option in RAxML) for each structural class and then performed a tree search in RAxML. We used the ML tree from Ryan et al. as a starting tree and estimated the model parameters. If the best tree obtained using the estimated parameters differed from the starting tree, we re-estimated model parameters using the new tree, iterating this procedure until the input and output tree converged (cf. Le and Gascuel 2010). Hereafter, the model parameters optimized for each structural class are further referred as structure-based model estimates; the parameter estimates are available from Zenodo (Pandey and Braun 2018).

We used multidimensional scaling in R (R Development Core Team 2011) to visualize the differences among estimates of exchange rate parameters obtained from the structural partitions as well as the standard empirical models. Since 20×20 exchangability matrix is symmetrical, we treated the half matrix of exchangability values for each model as a vector with 190 elements, normalized the elements to sum to one, and calculated the Euclidean distances among those vectors of exchange rate parameters. Then we used R cmdscale function to reduce this matrix to two dimensions (R script link: https://github.com/aakanksha12/Multidimensional-Scaling/blob/master/MDS_NoTM.md).

To test whether the different rate matrix parameter estimates might differ due to sampling variance alone we randomly sampled between 500 and 55,000 sites from the original dataset and estimated GTR exchangability parameters on those random samples. We then calculated the Euclidean distance between the GTR exchangability parameters estimated using the complete concatenated dataset (the grand GTR parameters) and those estimated using the randomly sampled subsets. These distances were then compared to the distance between GTR parameters estimated using each structural classes and the overall concatenated dataset GTR.

### Analyses using site heterogeneous models

To investigate the capability of site heterogeneous models to explain the differences between exposed and buried residues, we used the ML based version of CAT (Le et al. 2008) model integrated into IQ-TREE v. 1.5.3 (Nguyen et al. 2015) for various profile mixture classes (C10 – C60). We ran various profile mixture classes (C10 – C60) with exposed and buried alignments with (I+G4+FO) options in the program. Nodal support was assessed using ultrafast bootstrap (Minh et al. 2013) with 1000 replicates (-bb 1000).

## Acknowledgements

We are grateful to Rebecca Kimball, Joe Ryan, and Gavin Naylor for helpful comments on this manuscript and encouragement throughout the project.

## Supplementary Figures

**Supplementary Fig. S1.**
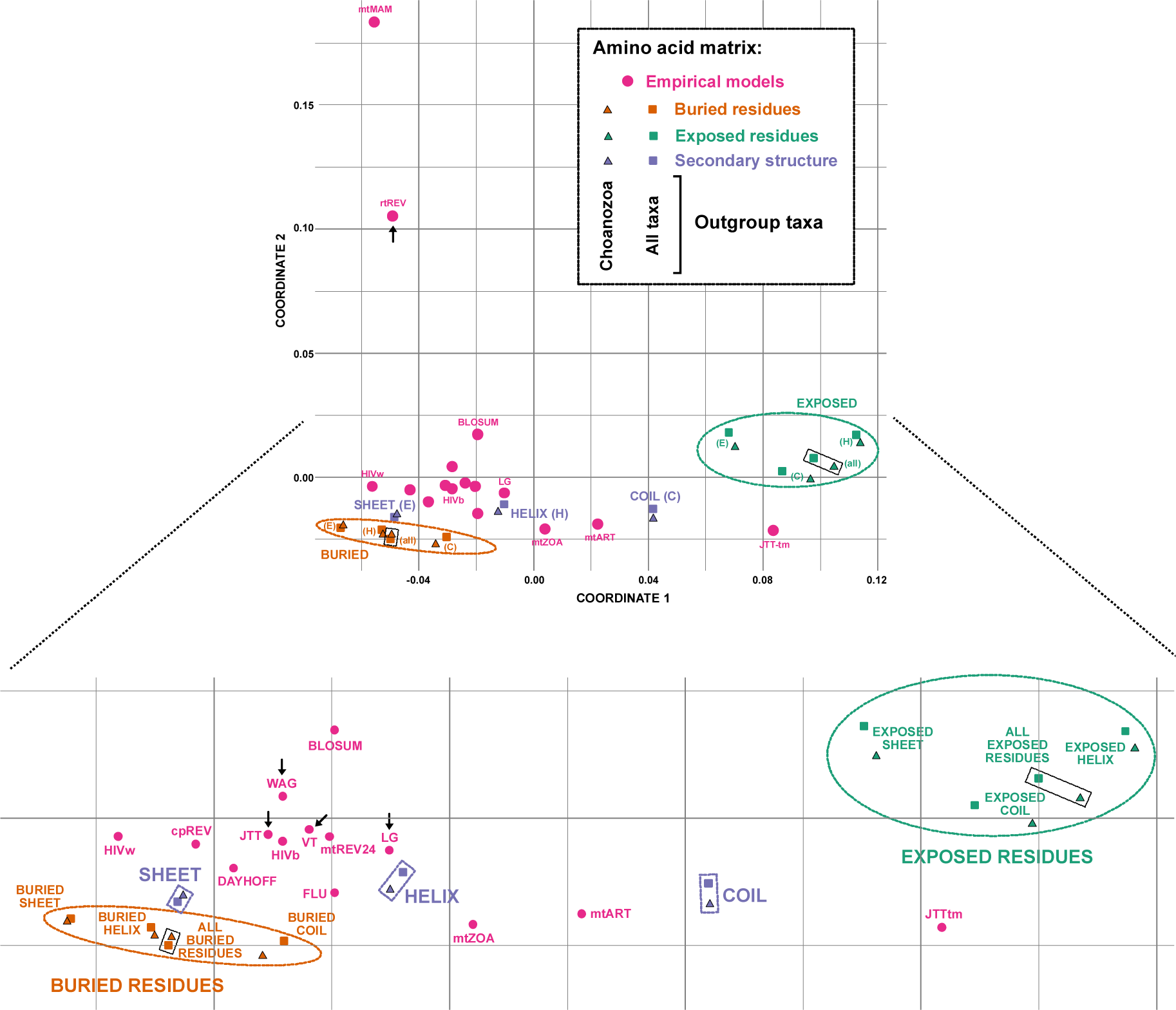
Multidimensional scaling plot based on Euclidean distances among amino acids exchange rate matrices. This plot expands fig. 4 to allow us to label all standard empirical models (pink circles); small arrows indicate the empirical models used in this study. Squares and triangles indicate rate matrices estimated using either all taxa (squares) or the reduced taxon set limited to choanozan outgroups (the Apoikozoa taxon set). Green and orange dotted ovals enclose clusters of rate matrices for exposed and buried residues, respectively; rate matrices based on all exposed or buried sites are emphasized by shaded boxes within those ovals. Rate matrices based on exposed or buried sites separated into subsets based on secondary structure are paired and labeled. Purple rectangular boxes indicate the three groups of rate matrices based on secondary structure (helix, sheet, and coil) without separating those residues into exposed and buried subsets.

**Supplementary Fig. S2.**
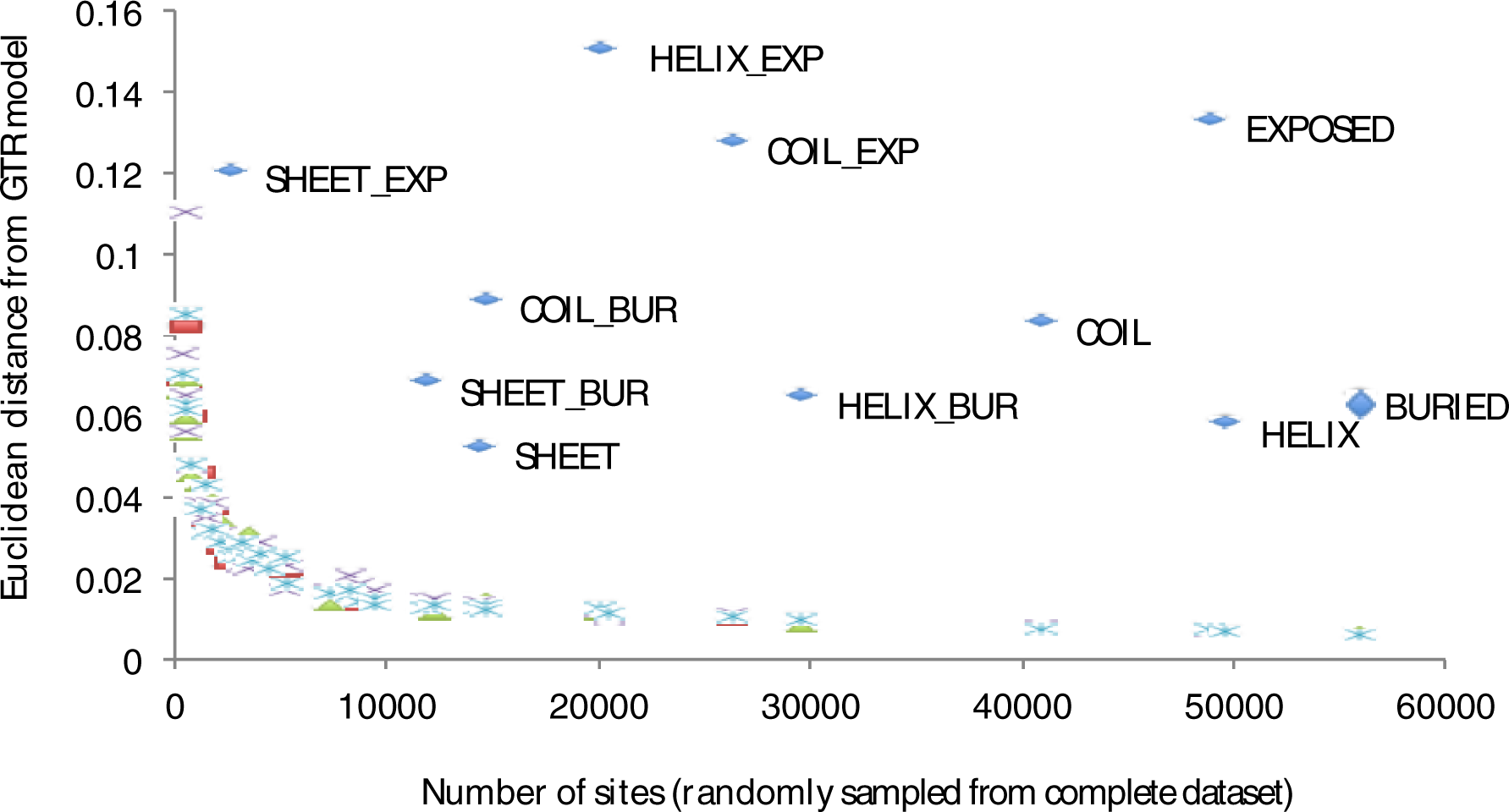
Euclidean distance between “grand” GTR rate matrix parameters (i.e., the rate matrix parameters estimated using the GTR model with the complete FRG dataset) and GTR rate matrix parameters estimated using subsets of the data. Rate matrix parameters optimized on sites defined using protein structure are presented as blue diamonds. The other points represent distances between the grand GTR model and rate matrix parameter estimates optimized using random samples (ranging in size from 500-55,000 aligned amino acid sites) that were drawn from the concatenated FRG dataset. For each data subset size, we generated 10 random samples; the distance between the rate matrix for structurally defined sites and the grand GTR model always exceeds the distance for random samples of sites of comparable size.

**Supplementary Fig. S3.**
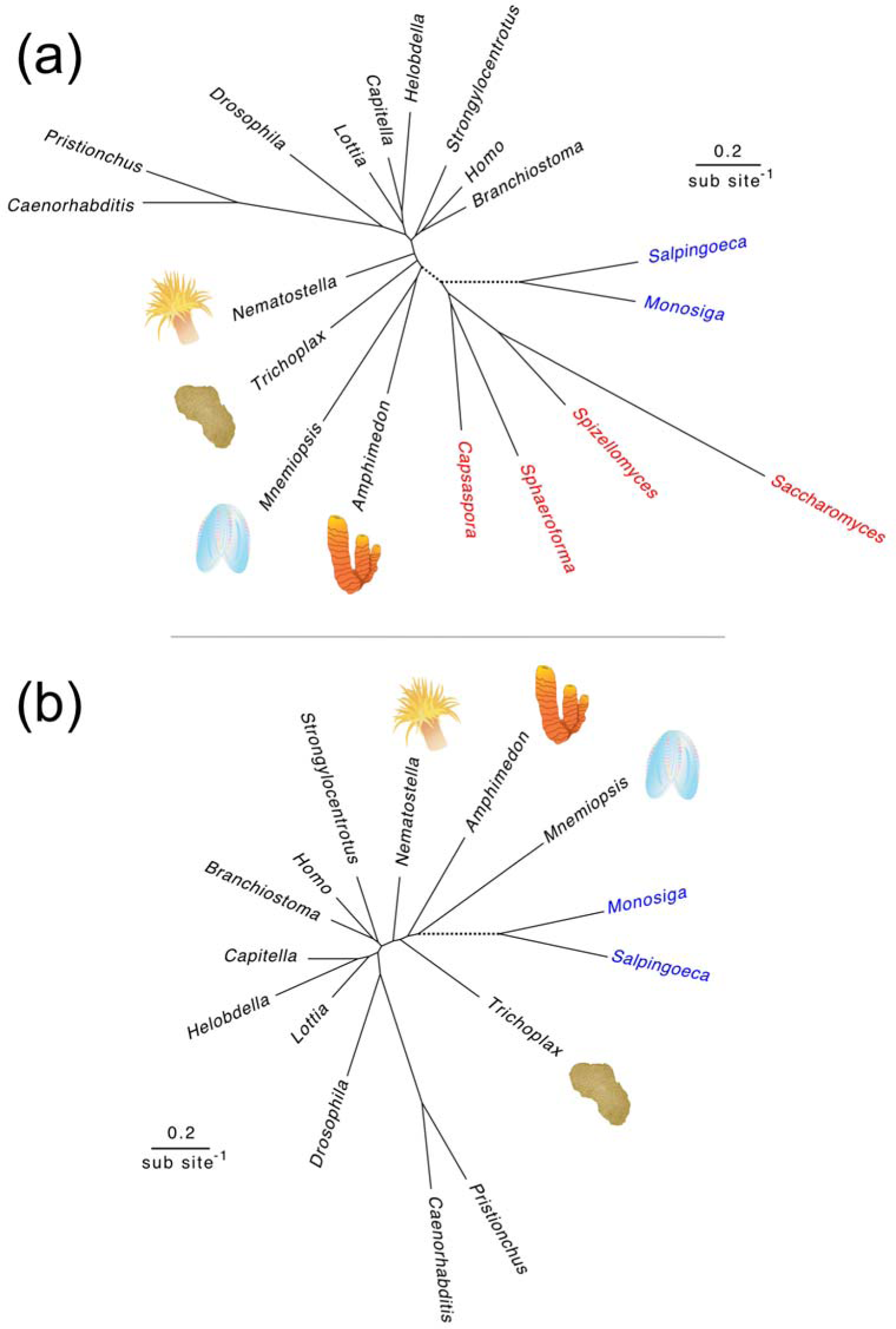
Estimates of metazoan phylogeny based on buried residues presented as unrooted phylograms. Outgroup taxa are emphasized using blue (Choanozoa) and red (other outgroups). (a) Complete dataset emphasizing. (b) Reduced (Apoikozoa only) taxon set. Note that the branch separating the choanozoan outgroups from the ingroups (indicated using a dashed line) is almost bisected by the other outgroup taxa. Drawings used to illustrate taxa are identical to those used to illustrate the major clades in fig. 1.

**Table S1.** Decisive sites obtained using T2 and T3 from fig. 2 with exposed and buried residues

|  | T2 | T3 |
| --- | --- | --- |
| Exposed | 248 | 151 |
| Buried | 228 | 255 |

T2 Tree obtained using EXPOSED residues with GTR optimized on exposed residues (fig. 2, left)
T3 Tree obtained from BURIED residues with GTR optimized on buried residues (fig. 2, right)

**Table S2.** Decisive sites obtained using T2* and T3* with Exposed and Buried residues

|  | T2* | T3* |
| --- | --- | --- |
| Exposed | 178 | 147 |
| Buried | 172 | 199 |

T2\* Tree with ctenophores sister to all other metazoans with (placozoa + cnidaria clade)
T3\* Tree with ctenophore + sponges clade tree (with placozoa + cnidaria clade)

